# Multi-‘Omic Integration via Similarity Network Fusion to Detect Molecular Subtypes of Aging

**DOI:** 10.1101/2022.11.16.516806

**Authors:** Mu Yang, Stuart Matan-Lithwick, Yanling Wang, Philip L De Jager, David A Bennett, Daniel Felsky

## Abstract

**Background:** Molecular subtyping of brain tissue provides insights into the heterogeneity of common neurodegenerative conditions, such as Alzheimer’s disease (AD). However, existing subtyping studies have mostly focused on single data modalities and only those individuals with severe cognitive impairment. To address these gaps, we applied Similarity Network Fusion (SNF), a method capable of integrating multiple high-dimensional multi-’omic data modalities simultaneously, to an elderly sample spanning the full spectrum of cognitive aging trajectories.

**Methods:** We analyzed human frontal cortex brain samples characterized by five ‘omic modalities: bulk RNA sequencing (18,629 genes), DNA methylation (53,932 cpg sites), histone H3K9 acetylation (26,384 peaks), proteomics (7,737 proteins), and metabolomics (654 metabolites). SNF followed by spectral clustering was used for subtype detection, and subtype numbers were determined by eigen-gap and rotation cost statistics. Normalized Mutual Information (NMI) determined the relative contribution of each modality to the fused network. Subtypes were characterized by associations with 13 age-related neuropathologies and cognitive decline.

**Results:** Fusion of all five data modalities (n=111) yielded two subtypes (n_S1_=53, n_S2_=58) which were nominally associated with diffuse amyloid plaques; however, this effect was not significant after correction for multiple testing. Histone acetylation (NMI=0.38), DNA methylation (NMI=0.18) and RNA abundance (NMI=0.15) contributed most strongly to this network. Secondary analysis integrating only these three modalities in a larger subsample (n=513) indicated support for both 3- and 5-subtype solutions, which had significant overlap, but showed varying degrees of internal stability and external validity. One subtype showed marked cognitive decline, which remained significant even after correcting for tests across both 3- and 5-subtype solutions (*p*_*Bonf*_=5.9×10^−3^). Comparison to single-modality subtypes demonstrated that the three-modal subtypes were able to uniquely capture cognitive variability. Comprehensive sensitivity analyses explored influences of sample size and cluster number parameters.

**Conclusion:** We identified highly integrative molecular subtypes of aging derived from multiple high dimensional, multi-’omic data modalities simultaneously. Fusing RNA abundance, DNA methylation, and H3K9 acetylation measures generated subtypes that were associated with cognitive decline. This work highlights the potential value and challenges of multi-’omic integration in unsupervised subtyping of postmortem brain.

## Introduction

Aging is often accompanied by progressive cognitive decline. The severity of this decline ranges from normal age-related changes to clinically important mild cognitive impairment (MCI) and ultimately dementia [1,2]. Alzheimer’s disease (AD) is the most common cause of late-life dementia, which is typically characterized by impairments in memory and loss of daily functioning [2]. This poses a major public health concern, as by 2050, the estimated number of individuals diagnosed with dementia globally is expected to reach 152.8 million [3]. As a neuropathological process, AD is defined by the abnormal accumulation of neurofibrillary tangles (hyperphosphorylated tau protein), the formation of extracellular dense core plaque deposits (beta-amyloid), and chronic neuroinflammation in the brain [4]. However, there is great inter-individual heterogeneity in these pathological hallmarks, and the relationship between neuropathology and cognitive impairment is not deterministic [5]. As such, there likely remain unobserved molecular signatures of age-related cognitive decline that could help explain the heterogeneity observed within populations and shed light on mechanisms of illness.

Molecular subtyping most often refers to classifying individuals within a population into subgroups using molecular data types and unsupervised clustering methods [6,7]. The approach has seen success in fields with abundant and readily assayed tissue samples from diseased populations, such as in oncology, where biopsied tumors yield molecular information leading to precision interventions [7]. Similarly, the heterogeneity of cognitive aging may be partly explained by using high-dimensional molecular measures from postmortem brain tissue of elderly donors to group similar individuals. For example, molecular subtypes of AD derived from RNA sequencing (RNAseq) data have been associated with AD-relevant pathologies [8–11], including amyloid and tau neuropathological burden, and *APOE* genotype [8,9]. Subtypes derived from common genetic variation, specifically single nucleotide polymorphisms, identified multiple AD-related molecular mechanisms [12]. A major limitation of most existing subtyping studies in this field is that they rely on information from single data modalities, e.g. gene expression data, which greatly constrains the information used to parse biological systems and pathological processes [13,14].

Importantly, it has been shown that several multi-’omic data types, including histone acetylation [15], metabolomics [16–20] and proteomics [21], are not only associated with AD neuropathologies, but also contributed information to associations that is missed with RNAseq alone [8,21]. As such, integrating data modalities into subtyping pipelines has been an active area of research [22,23], and large-scale cohort studies of aging that include brain donation and multi-‘omic characterization, such as those from the Accelerating Medicines Partnership for Alzheimer’s Disease (AMP-AD) consortium, now offer opportunities for developing highly integrative models of cognitive decline [24]. Methods development in high-dimensional feature integration have also facilitated these analyses [25,26], though not yet in pathological aging or AD. Similarity network fusion (SNF) is a network-based method specifically developed to integrate several multi-’omic data modalities simultaneously [27].

Here we performed a highly integrative analysis on up to five postmortem multi-’omic data modalities simultaneously, measured in the same individuals, to identify molecular subtypes of aging using the SNF method. We then characterized these subtypes by associating subtype membership with 13 age-related neuropathologies, antemortem cognitive performance, and rates of longitudinal cognitive decline. The most important features contributing to the fully fused similarity network were identified and subsequent analyses focused on the most informative data modalities. Lastly, we performed comprehensive sensitivity testing to explore the effects of parameter selection in unsupervised multi-’omic subtyping, which are often chosen arbitrarily.

## Methods

### Study participants

Data were analyzed from two longitudinal cohort studies of aging and dementia: the Religious Orders Study and Rush Memory and Aging Project (ROS/MAP), with more than 3,500 predominantly white elderly (mean = 78.44, sd = 7.79) participants of mostly European descent without known dementia at the time of enrollment [28]. Participants in ROS (1994-present) are older Catholic priests, nuns, and brothers across the United States, whereas MAP (1997-ongoing) recruits primarily from retirement communities and via social service agencies and Church groups throughout northeastern Illinois [28,29]. Combined data analysis for these two cohorts are enabled by harmonized protocols for participant recruitment, clinical assessment, and neuropathological examination at autopsy (autopsy rate exceeding 86%) with a large common core of identical item level data. A Rush University Medical Center Institutional Review Board approved each study. All participants signed an Anatomic Gift Act as well as informed and repository consents. Annual visits include tests of cognition function and a broad range of other demographic, social, lifestyle, and clinical assessments with an averaged follow-up rate of 97% [29]. Further details about the ROS and MAP cohorts can be found in previous publications [30] and through the Rush Alzheimer’s Disease Center Research Resource Sharing Hub, where participant-level clinical and demographic data are available via restricted access (https://www.radc.rush.edu/home.htm).

### Multi-’omic data used for subtyping

We used five multi-’omic data modalities to identify molecular subtypes: bulk RNAseq (18,629 genes, n_RNAseq_=1,092), DNA methylation (53,932 cpg sites, n_DNA_=740), histone H3K9 acetylation (26,384 peaks, n_histone_=669), metabolomics (654 metabolites, n_metabolomics_=514), and tandem mass tag (TMT) proteomics (7,737 proteins, n_proteomics_=368). All data types were acquired from the same brain region postmortem: dorsolateral prefrontal cortex (DLPFC). All ‘omic datasets used in our analyses were generated by members of the Accelerating Medicines Partnership - Alzheimer’s disease (AMP-AD) consortium and are available via restricted access through the AMP-AD knowledge portal, on Synapse (https://adknowledgeportal.synapse.org/). Further details can be found in Acknowledgements.

### RNA sequencing (RNAseq)

Full details on gene-level expression data from bulk DLPFC tissue have been published [31]. Approximately 100 mg of DLPFC tissue were dissected from autopsied brains. Samples were processed in batches of 12–24 samples for RNA extraction using the Qiagen MiRNeasy Mini (cat no. 217004) protocol, including the optional DNAse digestion step. RNA Samples were submitted to the Broad Institute’s Genomics Platform for transcriptome library construction following sequencing in three batches using the Illumina HiSeq (batch #1: 50M 101bp paired end reads) and NovaSeq6000 (batch #2: 30M 100bp paired end; batch#3: 40-50M 150bp paired end 121 reads) [32]. A cut-off point of 5 for RNA Integrity Number (RIN) score was used for constructing the cDNA library [33]. The average sequencing depth was 50 million paired reads per sample. To achieve higher quality of alignment results, a paralleled and automatic RNAseq pipeline was implemented based on several Picard metrics (http://broadinstitute.github.io/picard/). 18,629 features - full-length gene transcripts - from 1,092 samples remained after data preprocessing and quality control (QC).

### DNA methylation

Tissues were dissected similar to gene-expression data, full details on DNA methylation data have been published [33]. DNA was extracted by the Qiagen QIAamp mini protocol (Part number 51306). Probes with p-value >0.01 were removed at probe level QC if predicted to cross-hybridize with sex chromosomes and having overlaps with known SNP with MAF ≥0.01 (±10 bp) based on the 1000 Genomes database. Subject level QC methods including principal component analysis and bisulfite conversion efficiency. β-values reported by the Illumina platform were used as the measurement of methylation level for each CpG probe tagged on the chip; where missing values were imputed by the k-nearest neighbor algorithm (k=100). The primary data analysis was adjusted by age, sex, and experiment batch [33]. Due to the large number of features present for this data type, and to limit computational time, we only included the top 53,932 methylation peaks showing the greatest variability (**Supplementary Figure 1A**). To verify that this selection process did not impact our subtyping efforts, we performed sensitivity analysis for 5-modal integration using all CpG sites - resulting subtype memberships were nearly identical (**Supplementary Figure 1B**).

### Histone H3K9 acetylation

For the acetylation of the ninth lysine of histone 3 (H3K9ac), which is a marker of open chromatin, the Millipore anti-H3K9ac mAb (catalog #06-942, lot: 31636) was identified as a robust monoclonal antibody for the chromatin immunoprecipitation experiment. Similar to RNAseq and DNA methylation, 50 milligrams of gray matter was dissected on ice from biopsies of the DLPFC of each participant of ROS/MAP. Chromatin labeled with the H3K9ac mark and bound to the antibody was purified with protein A Sepharose beads [15]. To quantify histone acetylation, single-end reads were aligned to the GRCh37 reference genome by the BWA algorithm after sequencing. Picard tools were used to flag duplicate reads. A combination of five ChIP-seq quality measures were employed to detect low quality samples: samples that did not reach (i) ≥15×106 uniquely mapped unique reads, (ii) non-redundant fraction≥0.3, (iii) cross correlation≥0.03, (iv) fraction of reads in peaks≥0.05 and (v) ≥6000 peaks were removed [15]. Samples passing QC were used to define a common set of peaks termed H3K9ac domains. H3K9ac domains of less than 100bp width were removed resulting in a total of 26,384 H3K9ac domains with a median width of 2,829 bp available for 669 subjects. Full details on H3K9ac data can be found on Synapse (https://www.synapse.org/#!Synapse:syn4896408).

### Metabolomics

Metabolomics data were generated by the Alzheimer’s Disease Metabolomics Consortium (ADMC; ADMC members list https://sites.duke.edu/adnimetab/team/), led by Dr. Rima Kaddurah-Daouk [18–20]. Metabolomic profiling of postmortem brain was conducted at Metabolon (Durham, NC) with the Discovery HD4 platform consisting of four independent ultra-high-performance liquid chromatography–tandem mass spectrometry (UPLC–MS/MS) instruments [16,17]. For the purpose of QC and better understanding of the underlying biological mechanisms, missing rates less than 20% on known metabolites and 40% on individuals were imposed. As SNF cannot handle missing data, random forest imputation [34] was then applied, resulting in 654 metabolites and 514 individuals. Full details on metabolomic assays and data processing can be found here (https://www.synapse.org/#!Synapse:syn26007830). The full metabolomics dataset and metadata can be accessed via the AMP-AD Knowledge Portal.

### Proteomics

Prior to TMT labeling, samples were randomized by co-variates (age, sex, postmortem interval (PMI), diagnosis, etc.), into 50 total batches (8 samples per batch) [35]. Peptides from each individual (n=400) and the GIS pooled standard (n=100) were labeled using the TMT 10-plex kit (ThermoFisher 90406). Peptide eluents were separated on a self-packed C18 (1.9 μm, Dr. Maisch) fused silica column (25 cm × 75 μM internal diameter) by a Dionex UltiMate 3000 RSLCnano liquid chromatography system (Thermo Fisher Scientific) [35,36]. Peptides were monitored on an Orbitrap Fusion mass spectrometer (Thermo Fisher Scientific). The mass spectrometer was set to acquire data in positive ion mode using data-dependent acquisition. Dynamic exclusion was set to exclude previously sequenced peaks for 20 s within a 10-ppm isolation window [35,36]. In this study we only include peptides and participants with a missing rate less than 20% followed by random forest imputation [37], resulting in 7,737 proteins and 386 individuals. Full details on proteomics data acquisition and processing can be found on synapse (https://www.synapse.org/#!Synapse:syn17015098).

#### Uniform multi-’omic feature post-processing

Due to differences in data feature preprocessing among the five selected ‘omic data modalities, we performed additional post-processing QC to determine whether technical and demographic covariates may be influencing global patterns of variability for each modality. To achieve this, we tested associations between age of death, sex, PMI, and study cohort (ROS vs. MAP) with each of the top 20 components from PCA for each ‘omic modality separately, as in previous ‘omic work in this cohort [31]. The proportion of variance explained by each PC from each of the five data modalities, and the corresponding associations of each PC with potential covariates, are shown in **Supplementary Figures 2-6**. Based on this assessment, we determined that four out of five data modalities showed significant associations of all four covariates within the first 10 principal components (RNAseq data had been post-processed already and residualized for each of these covariates in addition to modality-specific confounders). We therefore proceeded by residualizing all features from each modality according to a linear model including all four covariates. This conservative approach ensured that contributions of each modality to latent subgroups were not unbalanced by different representations of covariate-specific effects. We also performed iterations of the analysis without correction, finding very similar but not identical subgroup memberships for 5-modal integration (**Supplementary Figure 7**).

#### Neuropathological assessment

All selected postmortem neuropathological variables analyzed in this study have been previously published in detail [29,38]. In addition to the outcome of NIA-Reagan neuropathological diagnosis of Alzheimer’s disease [5,39], we examined 13 other individual pathologies: brainwide amyloid-beta, diffuse and neuritic plaque counts, paired helical filament tau, neurofibrillary tangle count, TDP-43 proteinopathy stage (4 levels), large vessel cerebral atherosclerosis rating (4 levels), arteriolosclerosis, semiquantitative summary of cerebral amyloid angiopathy pathology (CAA; 4 levels), pathologic stage of Lewy body disease (4 stages), gross chronic cerebral infarcts (coded as binary; presence/absence of infarcts), and cerebral microinfarcts (coded as binary; presence/absence of infarcts).

#### Cognitive performance and residual cognition (resilience)

Scores from five cognitive domains (episodic memory, semantic memory, working memory, perceptual speed and perceptual orientation) were recorded at last study visit and summarized by z-scoring for a composite measure of global cognition, as described [40]. In our study, we defined the last available global cognitive measure as cognitive performance proximal to death. Cognitive slopes were also derived from the same set of z-scores over time to measure the longitudinal aspect of cognitive decline [41]. To assess the resilience component of an individual’s cognitive capacity, we used the residual cognition approach [42,43]. Residual cognition was defined as the residuals of a linear model of global cognitive performance at last visit regressed on observed neuropathologies (beta-amyloid, neurofibrillary tangles, neuritic plaques, diffuse plaques, Lewy bodies, macroscopic infarcts, microscopic infarcts, atherosclerosis, arteriolosclerosis, TDP-43 and CAA).

### Statistical Analysis

#### Subtype identification with Similarity Network Fusion (SNF)

The Similarity Network Fusion (SNF) method was used to integrate multi-’omic data modalities [27]. SNF first constructs sample-by-sample similarity matrices for each data modality separately and then iteratively updates and integrates these matrices via nonlinear combination until convergence is reached, generating a fused similarity network [44]. SNF does not require any prior feature selection, but fully imputed (non-missing) data is required. According to best practices [35,37], random forest imputation was applied on both metabolomics and proteomics data to impute missing values. The ‘SNFtool’ R package (v2.2.0) was used for the network fusion pipeline, with recommended parameters K=40, alpha=0.5, and T=50 (where K is the number of neighbors used to construct the similarity matrices; alpha is a hyper-parameter used in the scaling of edge weights; T is the total number of algorithmic iterations). Spectral clustering, an unsupervised soft clustering method rooted in graph theory [27,45], is the default clustering method for ‘SNFtool’; it was applied to the full fused affinity matrix to cluster study participants into subtypes. Optimal cluster numbers were identified (2 to 8 clusters) by the rotation cost [46] and eigen-gap [45] methods. Data modalities contributing the most information to fused similarity matrices were computed by Normalized Mutual Information (NMI). NMI is a measure of relevance and redundancy among features [47], which helps to identify the data types that contribute most strongly to the fused network estimated by SNF [27].

#### Assessment of internal subtype validity

Due to the high dimensionality and heterogeneity of multi-’omic data, assessments of cluster validity are critical to tackling potential biases of clustering algorithms toward particular cluster properties and to evaluate the probability that clusters do in fact exist [48,49]. Upon subtype identification, we conducted internal cluster stability analysis using the R package ‘clValid’, which measures cluster validity and stability through several metrics derived from resampling and cross-validation. Metrics included in our studies are the average proportion of non-overlap (APN) and the average distance between means (ADM), which work especially well if the data are highly correlated, which is often the case in high-throughput genomic data [49–51]. For resampling, we pulled 80% of participants for a total of 300 random draws, in accordance with previously published work using the SNF pipeline [52] as well as other AD molecular subtyping efforts [8]. The adjusted Rand index (ARI) was used to measure the agreement between subtype membership solutions (ranging from 0 to 1, where ARI = 1 indicating perfect agreement) [53]. Chi-square statistics were also used to compare the independence between different subtyping solutions [54].

#### Identifying top individual features defining molecular subtypes

In order to identify molecular features that differed most between subtypes after spectral clustering, we performed one-way ANOVA tests between each normalized feature from each multi-’omic data modality and subtype groupings. P-values from F-tests were used as the measure of significance to rank features from each modality. Gene annotations for DNA methylation data were mapped using the UCSC genome browser [55], and histone acetylation peaks were annotated by Klein et al. [15].

#### Association of subtypes with neuropathology, cognition, and residual cognition

For each clustering solution, subtype membership was initially characterized by associations with 13 neuropathologies and three cognitive measures described above using linear or logistic regression. Subtype membership for each participant was represented with dummy variables for inclusion in each model (n_subtypes_-1). For models of neuropathology, co-variates included age at death, biological sex, educational attainment (years), PMI, study cohort, and *APOE* ε4 status.

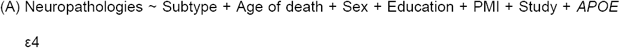

When fitting regression models for cognitive outcomes, the model (B) was also adjusted for the measurement latency, which is equal to the time difference (in years) between the last study visit where cognitive performance was assessed and age of death.

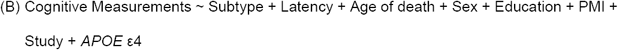

Omnibus F-tests of the hypothesis of equal outcome means (or probabilities for logistic models) across all subtypes were used to test the significance of subtype membership effects. *P*-values were Bonferroni adjusted for 16 tested outcomes, except where otherwise indicated. For subtypes with significant effects on global cognition (either at last visit or longitudinal slope), secondary analyses were performed (according to model B) for each subdomain of cognition separately.

### Sensitivity analyses for external validity across data modalities, sample sizes, and cluster numbers

To better understand the added value of data integration in the context of molecular subtyping, we performed a set of sensitivity analyses to measure differences in neuropathological and cognitive relevance (external validity) of subtypes derived from different combinations of multi-‘omic data modalities. Given that each iteration of these integrative analyses was limited to the sample size in which all data types were non-missing, we also assessed the effects of performing clustering in artificially limited sample subsets (i.e., where included non-missing data modalities permit a larger sample size). To achieve this, we defined a full search space of analytical pipeline configuration and parameter combinations for exhaustive modeling: 1) data modalities included (*d*; 31 possible combinations), 2) sample size (*n*; ranging from 111 to 1,092 participants, including 31 possible sample sizes each corresponding to a different data modality combination), and 3) cluster number (*c;* ranging from 2-5, the extremes of values observed in our subtyping analyses). This resulted in a total of 844 unique combinations of *d, n*, and *c*. To evaluate external validity, we performed omnibus tests of the association of subtype membership for each analytical iteration with the set of neuropathologies and cognitive measures, as previously. To provide some generalized insight into the effects of manipulating design parameters on our association strengths, second-level analyses were performed by relating each pipeline parameter to observed omnibus model significance for each neuropathology and cognitive outcome (*j*). For these analyses, the effects of *c* (and *cf*, the same parameter but treated as a categorical variable), *n*, and a new parameter, *m*, representing the number of data modalities being fused, were tested independently, according to the following formulae:

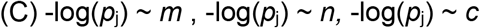

## Results

We analyzed data from a total of 1,314 unique participants from the Religious Orders Study and Memory and Aging Project (ROS/MAP) with at least one available multi-’omic data modality and non-missing clinical and neuropathological data. Sample demographics are summarized in **Table 1**. The number of participants with different degrees of overlapping multi-’omic characteristics ranged from n=111 (all five data types) to 1,092 (RNAseq only); all overlaps are shown in **Figure 1A**.

**Table 1.**
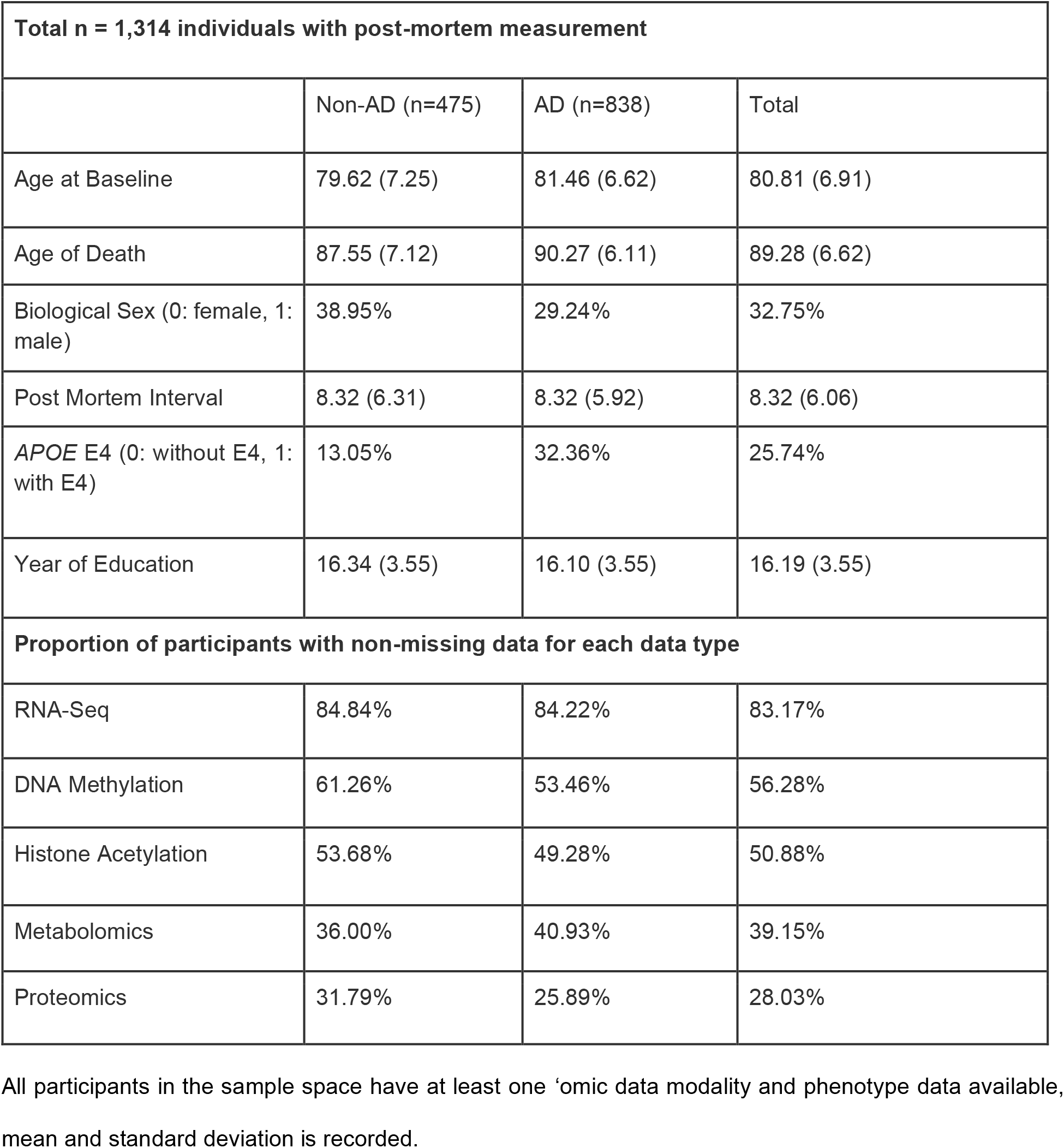
Table summarizing demographic data and the availability of multi-’omic data modalities stratified by NIA-Reagan diagnosis criteria in ROS/MAP.

| Total n = 1,314 individuals with post-mortem measurement |  |  |  |
| --- | --- | --- | --- |
|  | Non-AD (n=475) | AD (n=838) | Total |
| Age at Baseline | 79.62 (7.25) | 81.46 (6.62) | 80.81 (6.91) |
| Age of Death | 87.55 (7.12) | 90.27 (6.11) | 89.28 (6.62) |
| Biological Sex (0: female, 1: male) | 38.95% | 29.24% | 32.75% |
| Post Mortem Interval | 8.32 (6.31) | 8.32 (5.92) | 8.32 (6.06) |
| APOE E4 (0: without E4, 1: with E4) | 13.05% | 32.36% | 25.74% |
| Year of Education | 16.34 (3.55) | 16.10 (3.55) | 16.19 (3.55) |
| Proportion of participants with non-missing data for each data type |  |  |  |
| RNA-Seq | 84.84% | 84.22% | 83.17% |
| DNA Methylation | 61.26% | 53.46% | 56.28% |
| Histone Acetylation | 53.68% | 49.28% | 50.88% |
| Metabolomics | 36.00% | 40.93% | 39.15% |
| Proteomics | 31.79% | 25.89% | 28.03% |

All participants in the sample space have at least one 'omic data modality and phenotype data available, mean and standard deviation is recorded.

**Figure 1.**
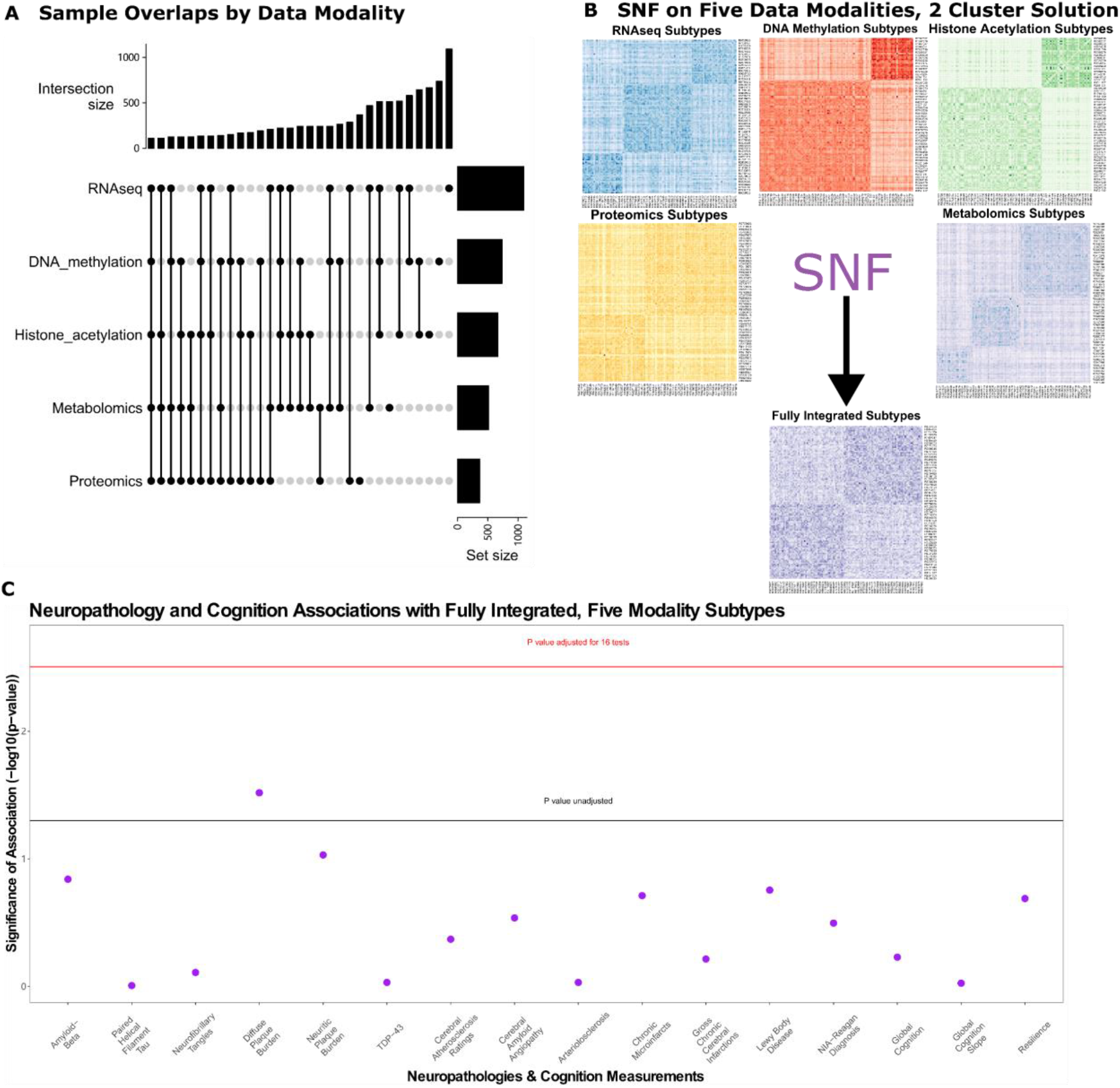
Molecular subtypes derived from 5 multi-’omic data modalities via SNF. A) Overlapping sample sizes across all combinations between five data modalities were examined using upset plot. B) Unimodal subtypes were identified from affinity matrices using spectral clustering accordingly from 111 overlapping samples (RNAseq: three subtypes, DNA methylation: two subtypes, histone acetylation: two subtypes, proteomics: two subtypes, metabolomics: three subtypes). Fully integrated subtypes were illustrated in the affinity matrix as well. C) Associations of fully integrated subtype memberships and 16 age-related neuropathologies and cognitive measurements were examined by omnibus F-tests for linear regression models. Y-axis shows significance of association (-log10 transformed raw p-values). The black horizontal line illustrates an unadjusted *p*-value threshold at 0.05, whereas the purple horizontal line demonstrates Bonferroni-adjusted *p*-value thresholds for 16 tests (*p*_raw_=3.1×10^−3^).

### Fully integrated five-modal network identifies two molecular subtypes nominally associated with neuritic plaque burden

First, we aimed to determine whether molecular subtypes derived from all five multi-’omic data modalities were informative of postmortem neuropathology and antemortem cognitive decline. SNF yielded an optimal solution of two molecular subtypes (**Figure 1B**) in 111 individuals with all five ‘omic modalities (n_S1_=53, n_S2_=58). Both the rotation cost and eigen-gap methods elected two as the optimal number of clusters. These subtypes were weakly associated with neuritic (*p*_raw_= 0.09) and diffuse plaque counts (*p*_raw_=0.03), though these associations did not survive correction for multiple testing. In addition, no significant associations were observed for cognitive performance at last visit, rate of cognitive decline, or residual cognition (**Figure 1C**).

Despite the lack of significant associations of molecular subtypes with pathology and cognition, the fully fused network demonstrated substantial internal stability (APN=8.7%, ADM=0.02; **Supplementary Figure 8**). We therefore proceeded to identify the data modalities contributing most to the fused network by normalized mutual information (NMI) (**Supplementary Table 1**). We found that histone acetylation (NMI=0.38), DNA methylation (NMI=0.18) and RNAseq (NMI=0.15) were the top contributors to the fused network (to a substantially greater degree than proteomic (NMI=0.04) and metabolomic modalities (NMI=0.05)). The top 10 individual features contributing to the fused network from the top contributors are summarized in **Supplementary Table 2**. Based on the importance of the top three data modalities, secondary analysis was conducted integrating only histone acetylation, DNA methylation, and RNAseq, which permitted subtyping of a much larger sample size with non-missing overlapping data (n=513).

### Subtypes derived from three-modal integration are associated with cognitive performance proximal to death and longitudinal cognitive decline

In secondary analyses with three data modalities, the eigen-gap method elected three molecular subtypes as the optimal clustering solution, while rotation cost elected five. We therefore evaluated both solutions by comparing membership overlap, differences in internal validity metrics, and associations with neuropathology and cognition. A strong overlap was identified between subtype memberships in the 3- and 5-subtype solutions (chi-square *p*=2.2×10^−16^, ARI=0.76; **Figure 2A, D**), whereby the large subtype 3 (n=377) from the 3-subtype solution contained 81.2% of the participants assigned to subtypes 3, 4, and 5 from the 5-subtype solution. Internal cluster stability was compared between 3-subtype and 5-subtype solutions (**Figure 2B, C**); both APN and ADM measures were better for the 3-subtype solution (APN=9.6%, ADM=0.01), though the 5-subtype solution also demonstrated cluster stability well above random chance (APN=23.1%, ADM=0.02) (**Figure 2E, F**).

**Figure 2.**
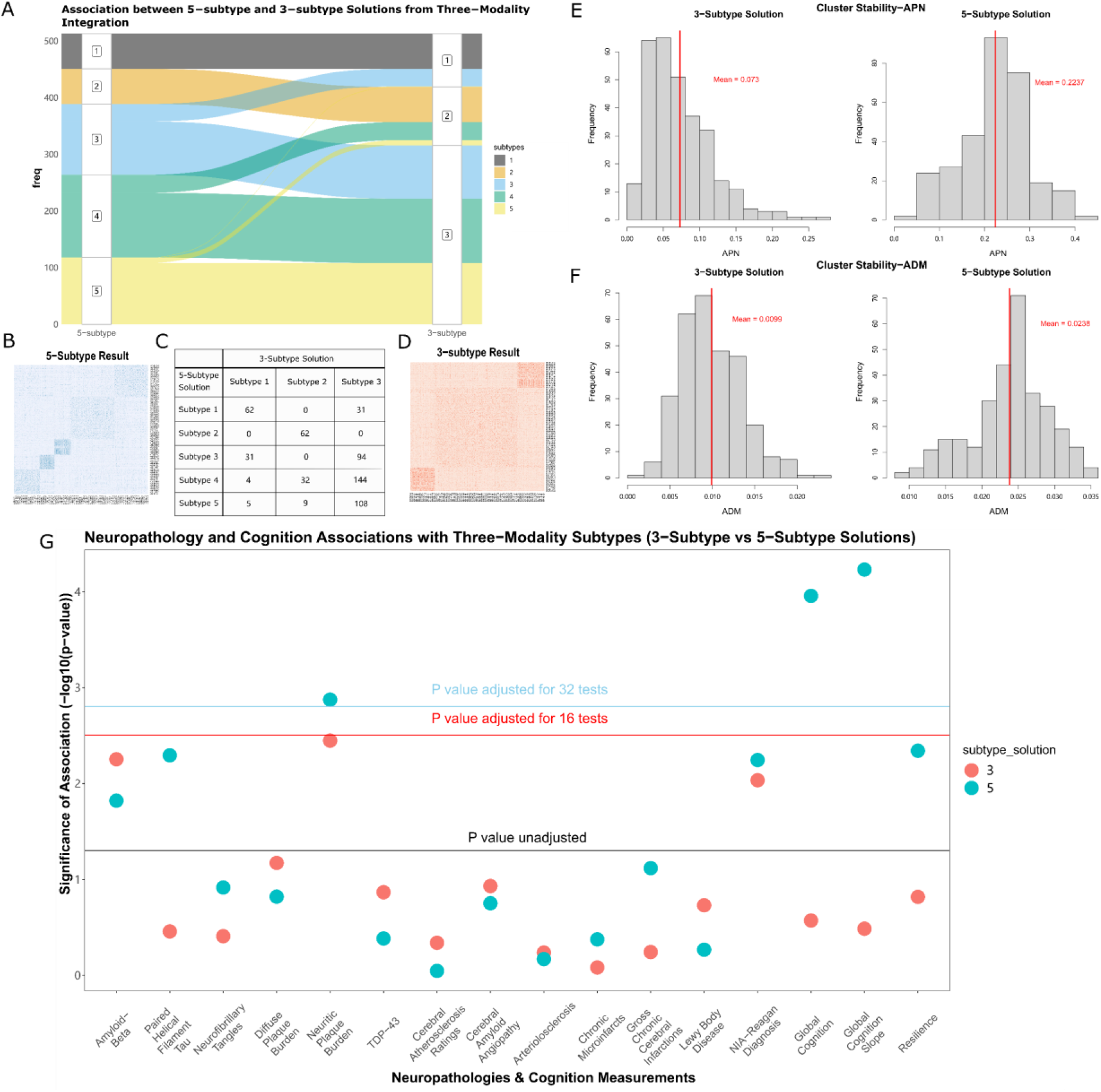
Two subtyping solutions derived from histone acetylation, DNA methylation and RNAseq were tested against each other both internally and externally. A) 3-subtype solution and 5-subtype solution derived from 3-modal integrated networks were associated with each other. B-D) Subtypes were identified from affinity matrices using spectral clustering, and overlapped with each other E) Histograms for the distribution of ADM generated from 300 random sub-samples for both 3-subtype and 5-subtype solutions. F) Histograms for the distribution of APN generated from 300 random sub-samples for both 3-subtype and 5-subtype solutions. G) Associations of 3-modal integrated memberships and 16 age-related neurobiological traits were examined by omnibus F-tests for linear regression models. Y-axis shows significance of association (-log10 transformed raw p-values). The black horizontal line illustrates an unadjusted *p*-value threshold at 0.05, whereas the red and blue horizontal lines demonstrate Bonferroni-adjusted *p*-value thresholds for 16 and 32 tests (*p*_raw_=3.1×10^−3^ and *p*_raw_=1.6×10^−3^), respectively. Two subtyping solutions for molecular subtyping were differentiated by color.

In tests of external validity, and tests of association with neuropathological and cognitive measures, subtype membership was significantly associated with global cognition at last visit (*p*_Bonf_=0.022) and rate of cognitive decline (*p*_Bonf_=4.2×10^−4^) for the 5-subtype solution after multiple testing correction (**Figure 2G**). In contrast, the 3-subtype solution was preferred by internal cluster stability metrics, and significant associations with neuropathology or cognition were not observed (**Figure 2G**). We therefore probed further into the 5-subtype solution.

Cross-tabulation of three-modal and five-modal subtype memberships was carried out for only the 111 individuals included in the full five-modal analysis above, finding substantial overlap (chi-square *p*=8.1×10^−9^, ARI=0.60; **Figure 3A**). This demonstrated that the SNF procedure was consistent across sample size in terms of defining core cluster memberships when the most influential data types were combined.

**Figure 3.**
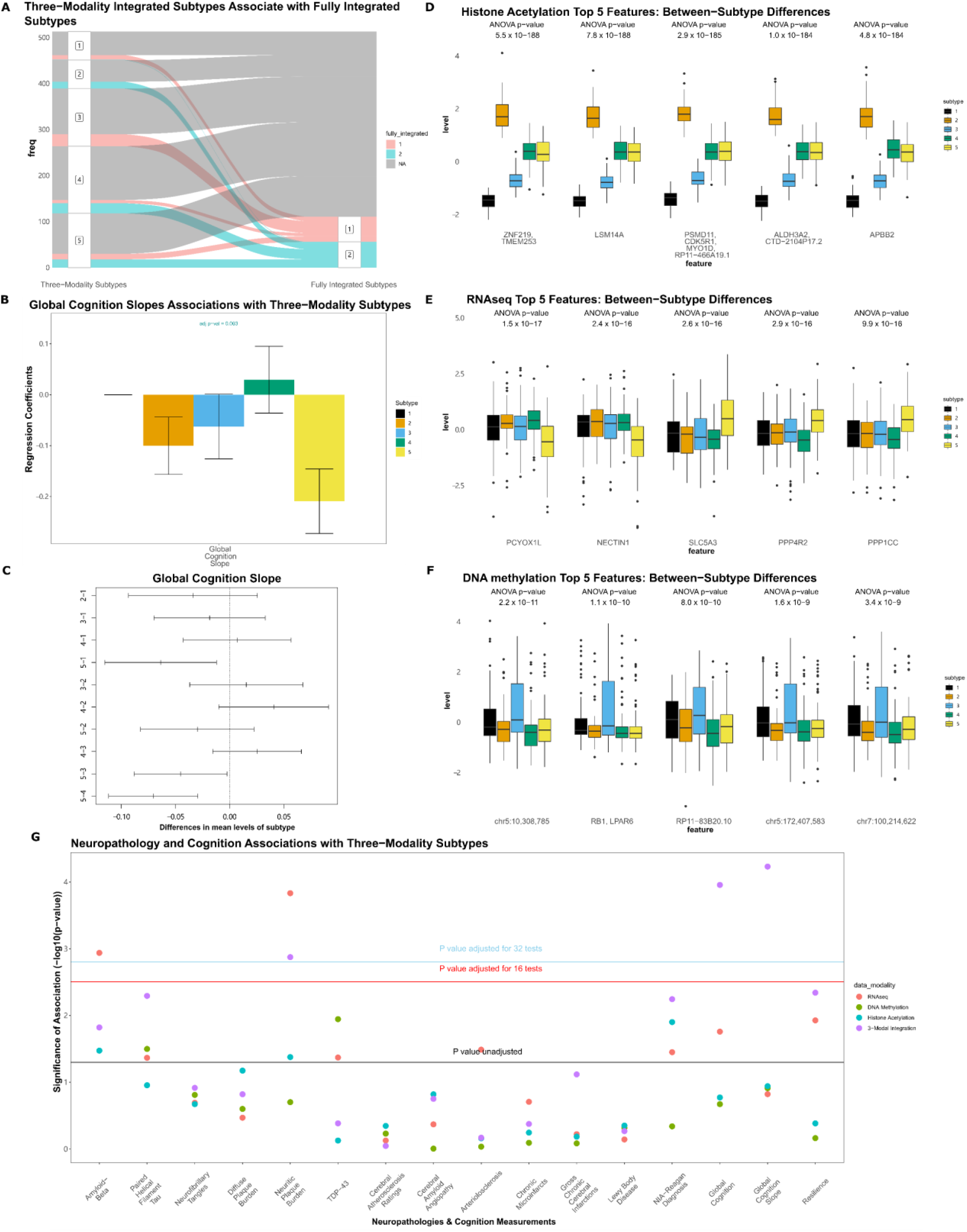
Molecular subtypes derived from histone acetylation, DNA methylation and RNAseq were tested against age-related neuropathologies and cognitive measurements. A) Subtypes derived from three-modal integrated networks were associated with the fully integrated subtypes. B) Consensus associations of three-modal integrated subtypes and rate of cognitive decline. Y-axis shows standardized beta coefficients estimated from linear regression, where subtype 1 was used as the baseline category (error bars show standard deviation from standardized linear regression models). C) Difference in mean value of rate of cognitive decline between subtypes by Tukey’s HSD. D-F) Boxplots showing the z-normalized values of the top 5 features contributing to the three-modal fused network from each input data modality. G) Associations of 3-modal integrated and unimodal subtype memberships with neuropathological and cognitive traits were examined by omnibus F-tests for linear regression models. Y-axis shows significance of association (-log10 transformed raw p-values). The black horizontal line illustrates an unadjusted *p*-value threshold at 0.05, whereas the red and blue horizontal lines demonstrate Bonferroni-adjusted *p*-value thresholds for 16 and 32 tests (*p*_raw_=3.1×10^−3^ and *p*_raw_=1.6×10^−3^), respectively. Data modalities used for molecular subtyping were differentiated by color.

In assessments of the mean differences in global cognition and the ratio of cognitive decline across 5 subtypes identified, subtype 5 had the worst global cognitive performance at last visit and the fastest rate of cognitive decline (**Figure 3B**). This difference was significant in post hoc pairwise tests against all other subtypes, except for subtype 2 (**Figure 3C**). Subtype 4 exhibited the best average cognitive performance and slowest decline (**Figure 3B, C**). Notably, the association observed with cognitive decline (*p*_Bonf_=5.9×10^−3^) was strong enough to survive correction for multiple testing across combined 5-subtype and 3-subtype association test sets (32 tests) (**Figure 2G**). Given the significant association of subtypes with global cognition at last visit and rate of global cognitive decline, we performed follow-up analysis on five cognitive subdomains. For rate of cognitive decline, subtypes were most strongly associated with perceptual orientation (*p*_Bonf_=8.0×10^−5^), perceptual speed (*p*_Bonf_=0.004), and semantic memory (*p*_Bonf_=0.007) (**Supplementary Figure 9A**). Specifically, the best and worst cognitive performance values were observed on average in subtypes 4 and 5, respectively (**Supplementary Figure 9B-F**). A similar pattern was also identified from cognition measured at last visit (**Supplementary Figure 9G-L**).

### Molecular features defining three-modal subtypes

To describe the molecular signals most strongly associated with our observed subtypes, we first identified the top features contributing to the fused network from each data modality by ANOVA (**Supplementary Table 3**). The top 5 histone acetylation features exhibited the strongest within-subtype homogeneity and between-subtype variability (consistent with the observation that histone acetylation had the largest NMI of each modality **Supplementary Table 1**). The most extreme values for acetylation were observed in subtypes 1 (lowest levels) and 2 (highest levels) at peaks annotated to *ZNF219, TMEM153, LSM14A, PSMD11, CDK5R1, MYD1D, ALDH3A2, APBB2*, and others (**Figure 3D**). Subtype 5, which was characterized by the fastest rate of cognitive decline, had intermediate acetylation of these peaks (along with subtype 4, which are largely represented by subtype 3 in the 3-subtype solution). For DNA methylation, CpG sites showed differential methylation at sites annotated to *RB1, LPAR6*, and *RP11-83B20*.*10*, as well as intergenic regions on chromosome 5 and 7, though no consistent pattern related to the cognition-associated subtype 5 was observed (**Figure 3F**). In contrast, the top subtype-associated RNAseq features revealed lower levels of *PCYOX1L* and *NECTIN1*, as well as higher levels of *SLC5A3, PPP4R2*, and *PPP1CC* in subtype 5 specifically compared to all other subtypes (**Figure 3E**).

### Comparison with single modality subtypes and sensitivity analysis

Finally, we compared clinical and neuropathological associations of these three-modal subtypes with those for subtypes derived from each of the modalities analyzed individually. We found that these integrated subtypes had unique associations with cognitive performance and decline. For example, subtypes derived from RNAseq alone (n=1,092) were significantly associated with amyloid-beta (*p*_Bonf_=0.018) and neuritic plaque burden (*p*_Bonf_ =2.3×10^−3^), but not with global cognition at last visit (*p*_Bonf_=0.28) or rate of cognitive decline (*p*_Bonf_=1.0). In fact, none of the unimodal subtypes showed more significant associations than three-modal, 5-cluster subtypes on global cognitive performance (**Figure 3G**).

In sensitivity analyses, substantial variability in external validity was observed across different selections of sample size, data modalities, and cluster number. **Supplementary Figure 10** illustrates the full set of results for selected amyloid and cognitive outcomes, which were the outcomes demonstrating the most significant associations with subtype membership in our analyses above (full summary statistics from these analyses are available in **Supplementary Table 4**). **Supplementary Figure 11A** shows the meta-regression results for the influence of sample size (*n*), number of data modalities (*m*), and cluster number (*c*) on statistical associations with all 16 tested phenotypes. Generally, less significant associations were captured as more data modalities were integrated and sample size decreased (see example of beta-amyloid in **Supplementary Figure 11B**), though exceptions were noted, such as for Lewy bodies (where additional modalities on average increased external validity; meta *p*_raw_=2.5×10^−7^; **Supplementary Figure 11C**). Comparatively, cluster number selection had less of an impact overall on external validity.

## Discussion

We used up to five ‘omic data modalities acquired from the human postmortem prefrontal cortex simultaneously to detect molecular subtypes of aging using a high-dimensional, unsupervised approach. We identified several subtypes that were significantly associated with individuals’ rates of cognitive decline and levels of beta-amyloid neuropathology. In particular, molecular subtypes derived from a three-modal integrated network combining gene expression (RNAseq), H3K9ac, and DNA methylation peaks yielded subtypes of participants with significantly faster decline in global cognition, specifically in domains of perceptual orientation, perceptual speed, and semantic memory. To the best of our knowledge, associations between multi-’omic subtypes and cognitive performance have not previously been identified, and most subtyping studies have focused only on individuals with confirmed, late-stage AD [56]. Our findings also empirically quantify the relative information provided by different ‘omic modalities to participant similarity networks.

In fully integrated analyses, combining all five available modalities, we identified two molecular subtypes which exhibited non-significant external validity with respect to neuropathology and cognition. We did not explore this result much further for three reasons: 1) both internal cluster validity metrics (eigen gap and rotation cost) elected the same 2-subtype solution, 2) the sample size for full five-modal integration analysis was small (n=111), and 3) NMI calculations showed substantial heterogeneity in the amount of information contained within each modality when considering patient similarity networks in this sample subset. The small sample size was likely a key limitation; this was confirmed by sensitivity analyses showing that even for single data modalities, when the sample was restricted to the n=111 group, there were virtually no observed associations with any cognitive or neuropathological measures.

By comparing both integrated molecular subtypes and unimodal subtypes from spectral clustering, we found that subtypes from RNAseq alone were significantly associated with neurofibrillary tangles and amyloid-beta. Such associations align with findings from previous subtyping work in only individuals suffering from dementia [8], and demonstrate the reliability of the method we used for subtyping. Our analysis also emphasizes the importance of integrating epigenetic data with gene expression studies seeking to identify key molecular drivers of AD [57]. Variability in gene expression alone cannot determine the current status of diseases [58,59]; even so, genetic and epigenetic studies still tend to be conducted separately [57]. This study serves as evidence that integrating multiple epigenetic data types with gene expression data can lead to the discovery of novel molecular subtypes associated with cognition.

In describing the top molecular features that distinguish our subtypes from one another, we identified epigenetic marks and RNA transcripts which map to genomic loci previously associated with AD and cognitive aging. Of particular interest were those loci that differentiated cognition-associated subtype 5 from all other subtypes. In this subtype, we found lower levels of Prenylcysteine Oxidase 1 Like (*PCYOX1L*), a gene which has been previously associated with AD [60–63], and has been identified as an AD target gene by the Agora platform (https://agora.adknowledgeportal.org/) with strong evidence for RNA down-regulation across 8 brain regions and proteomic down-regulation across four regions. Nectin cell adhesion molecule 1 (*NECTIN1*) [64] was similarly downregulated in subtype 5, and also showed RNA and protein-level dysregulation in the Agora database, confirming that the multi-modal SNF pipeline was capable of extracting some known signals with neuropathological significance.

Among the top genes with higher average levels in subtype 5 were *SLC5A3* [65], *PPP4R2*, and *PPP1CC. PPP4R2* and *PPP1CC* code for enzymes in the serine/threonine-protein phosphatase family and are well-known contributors to canonical AD pathological cascades [66]. Interestingly, *PPP4R2* has also been identified as a top hypomethylated gene of interest in a methylome-wide association study of Parkinson’s disease [67], an illness which is also often accompanied by cognitive decline [68]. Other top contributors to the three-modal subtypes, such as *PSMD11* [69], *APBB2* [70], and *TMEM253* [71] are also known to be involved in the development of AD pathology. *TMEM253* is also linked with mild cognitive impairment (MCI) via predicted gene expression based on genetic variation (TWAS) [71]. However, some top genes (e.g. *ZNF219*, a Kruppel-like zinc finger gene, has been associated with a-synucleinopathy [72] and has binding sites in the *MAPT* gene [73]). In contrast, these genes have not yet been associated with AD or cognitive aging, and our method provides a full resource of ranked importance for all ‘omic features studied, which provides novel targets for future study.

There are several limitations to consider when interpreting our results. First, a common challenge in unsupervised clustering endeavors, we did not achieve consensus on optimal clustering solutions in our three-modal subtyping analysis. In our case, we not only examined the optimal cluster number from two established methods especially suited to the SNF pipeline, but also tested cluster validity by multiple resampling measures, as there is no ground truth to compare to, and important information may be missed by heuristic methods alone [74],[75]. In our analysis, the disagreement between optimal cluster number as elected by internal stability measures vs. external cognitive and neuropathological information also demonstrates the importance of transparency in the presentation of clustering analyses; in our case, both the 3- and 5-subtype solutions had significant overlaps in identity, though only the fifth cluster revealed a significant cognitive deficit. We again emphasize that these effects on cognition would survive correction for multiple testing in a full pool of tests combining both 3- and 5-subtype solutions.

Second, differences in data preprocessing methods for our five ‘omic data modalities may have impacted downstream clustering, despite our efforts to control for technical and biological confounders at both the individual feature level and at the overall sample level in models testing external validity. Third, ROS/MAP is intrinsically limited by its inclusion of predominantly individuals of European-Caucasian ancestry, with an overrepresentation of biologically female participants [28–30]. Finally, ROS/MAP is known to be a resilient cohort of elderly individuals including some members of the religious communities of Illinois. Even though we modeled study as a covariate in all analyses to mitigate variability due to large lifestyle differences, results derived from such a study population might not be applicable to the entire population. Future studies will be required using populations with increased diversity with respect to ancestry and socio-demographics. This will be the means to achieve a better understanding of the degree to which our findings can be applied more broadly beyond European-Caucasians.

## List of Abbreviations

AD: late-onset Alzheimer’s disease
ADM: average distance between means
AMP-AD: Accelerating Medicines Partnership for Alzheimer’s Disease
APN: average proportion of non-overlap
ARI: adjusted Rand index
CAA: cerebral amyloid angiopathy
DLPFC: dorsolateral prefrontal cortex
H3K9ac: acetylation at the 9th lysine residue of the histone H3 protein
MAP: Rush Memory and Aging Project
MCI: mild cognitive impairment
NMI: Normalized Mutual Information
PCA: Principal component analysis
PMI: Post mortem interval
RNAseq: RNA sequencing
ROS: Religious Orders Study
SNF: Similarity Network Fusion
TMT: tandem mass tag

## Declarations

### Ethics approval and consent to participate

For The Religious Orders Study and Rush Memory and Aging Project, all study participants provided informed consent and both studies were approved by a Rush University Institutional Review Board. Further, all participants signed an Anatomic Gift Act for organ donation and signed a repository consent for resource sharing. For the Mayo dataset, protocols were approved by the Mayo Clinic Institutional Review Board and all subjects or next of kin provided informed consent.

### Consent for publication

Not applicable.

### Availability of data and materials

All multi-’omic datasets supporting the conclusions of this article are available via approved access at the Synapse AMP-AD Knowledge Portal (https://adknowledgeportal.synapse.org/, doi: 10.7303/syn2580853). All analyses were performed using open-source software. No custom algorithms or software were used that are central to the research or not yet described in published literature. ROSMAP resources can be requested at https://www.radc.rush.edu.

### Authors’ contributions

MY was responsible for data processing, statistical analysis, manuscript writing, and editing. SML contributed to manuscript editing. DF was responsible for data access, ensuring data quality control, study design, and manuscript writing and editing. YW, PLDJ and DAB were responsible for aspects of data collection, collaborative input on study design, and manuscript editing.

## Acknowledgements

The authors acknowledge all of the patients and their families for graciously donating brain tissue. ROS/MAP is supported by Yanling Wang, Philip L De Jager and David A Bennett. Daniel Felsky is supported by the Michael and Sonja Koerner New Scientist Award, the Krembil Family Foundation, the Canadian Institutes of Health Research, and the CAMH Discovery Fund. ROS/MAP is supported by NIH grants P30AG72971, P30AG72975, R01AG15819, R01AG17917, and U01jAG61356. ROS/MAP resources can be requested at www.radc.rush.edu. Results published here are based on metabolomics data generated by the Alzheimer’s Disease Metabolomics Consortium (ADMC; ADMC members list https://sites.duke.edu/adnimetab/team/), led by Dr. Rima Kaddurah-Daouk at Duke University, using biospecimens provided by the Rush Alzheimer’s Disease Center, Rush University Medical Center, Chicago. Support for the biospecimen processing and data generation conducted by the ADMC was provided by the following National Institute on Aging grants R01AG046171, RF1AG051550, RF1AG057452, RF1AG059093, RF1AG058942, U01AG061359, and the Foundation for the NIH (FNIH) grant: #DAOU16AMPA.

