## Supplementary Materials for "Multi-‘Omic Integration via Similarity Network Fusion to Detect Molecular Subtypes of Aging"

### Supplementary Figures

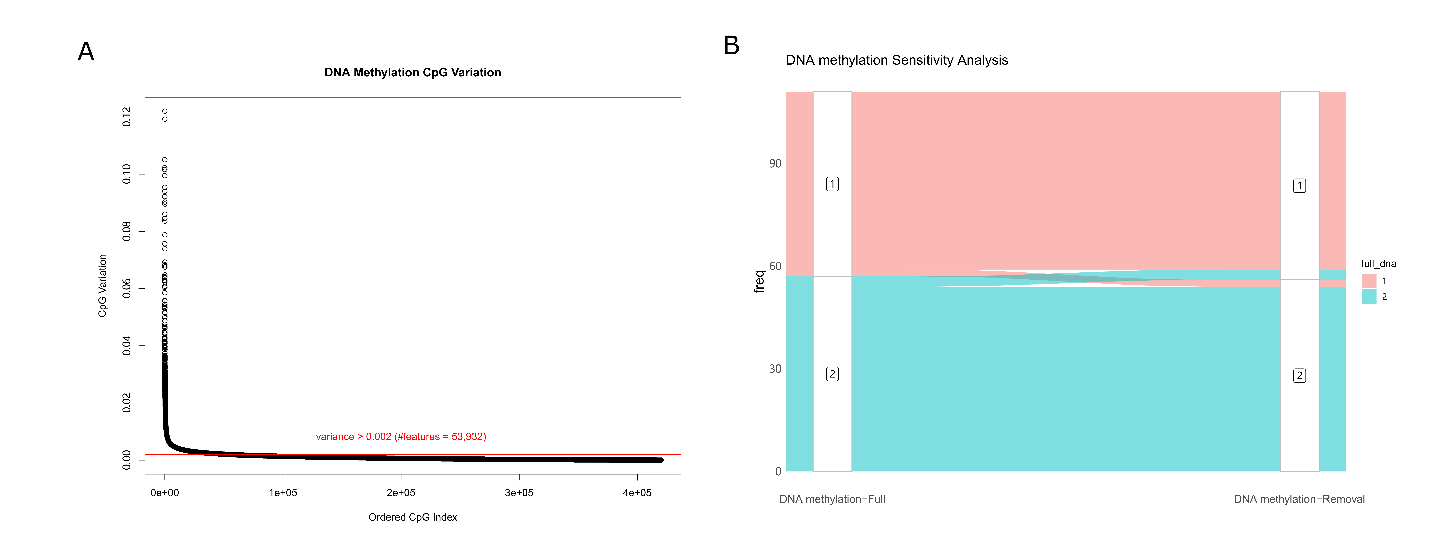

###### *Supplementary Figure 1. 361,916 CpG sites were removed due to low variation and the sensitivity test A) Ranked CpG variation across DNA methylation data. Y-axis describes the variation of each CpG site while x-axis describes the rank of CpG variation. The red horizontal line labels the threshold (> 0.002) of removal keeping 53.932 CpG sites. B) Alluvial plot comparing the subtype results on individual level before and after CpG sites removal. The optimal subtype number was assigned as 2.*

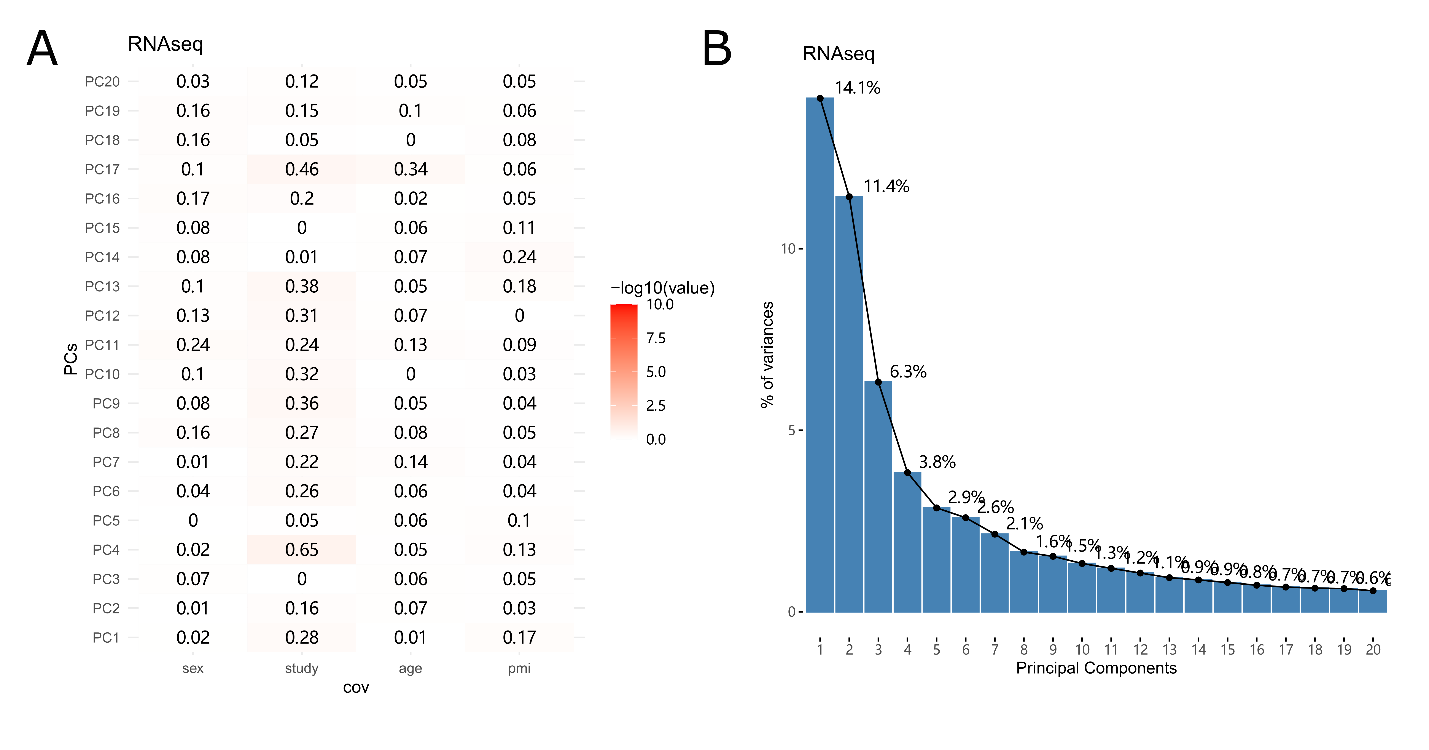

###### *Supplementary Figure 2. Top 20 Principal Components (PCs) from RNAseq data A) Significance of associations between top 20 PCs and four covariates derived from RNAseq data, including sex, study, age of death and PMI, derived from Omnibus F-tests for linear regression models B) Scree plots illustrating variances explained by top 20 PCs for RNAseq data.*

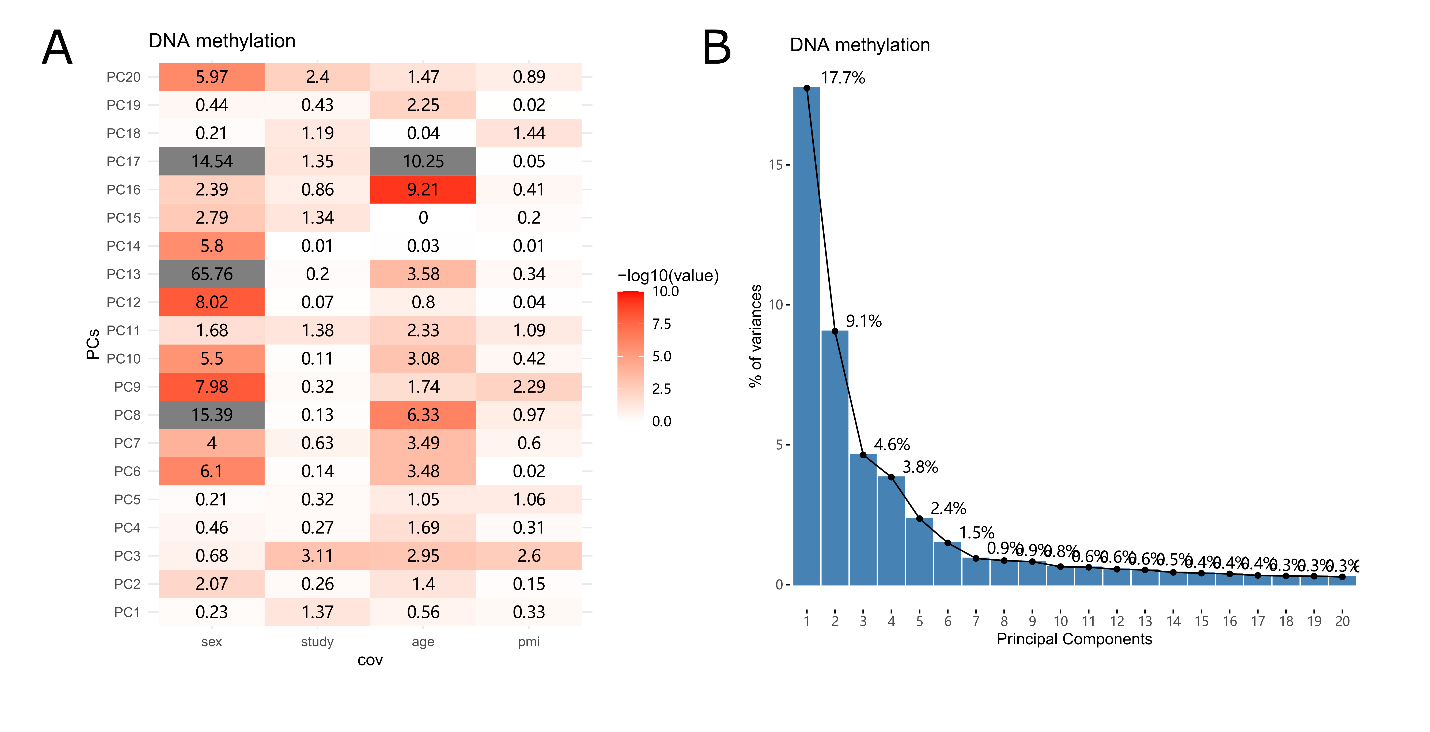

###### *Supplementary Figure 3. Top 20 Principal Components (PCs) from DNA methylation data A) Significance of associations between top 20 PCs and four covariates derived from DNA methylation data, including sex, study, age of death and PMI, derived from Omnibus F-tests for linear regression models B) Scree plots illustrating variances explained by top 20 PCs for DNA methylation data.*

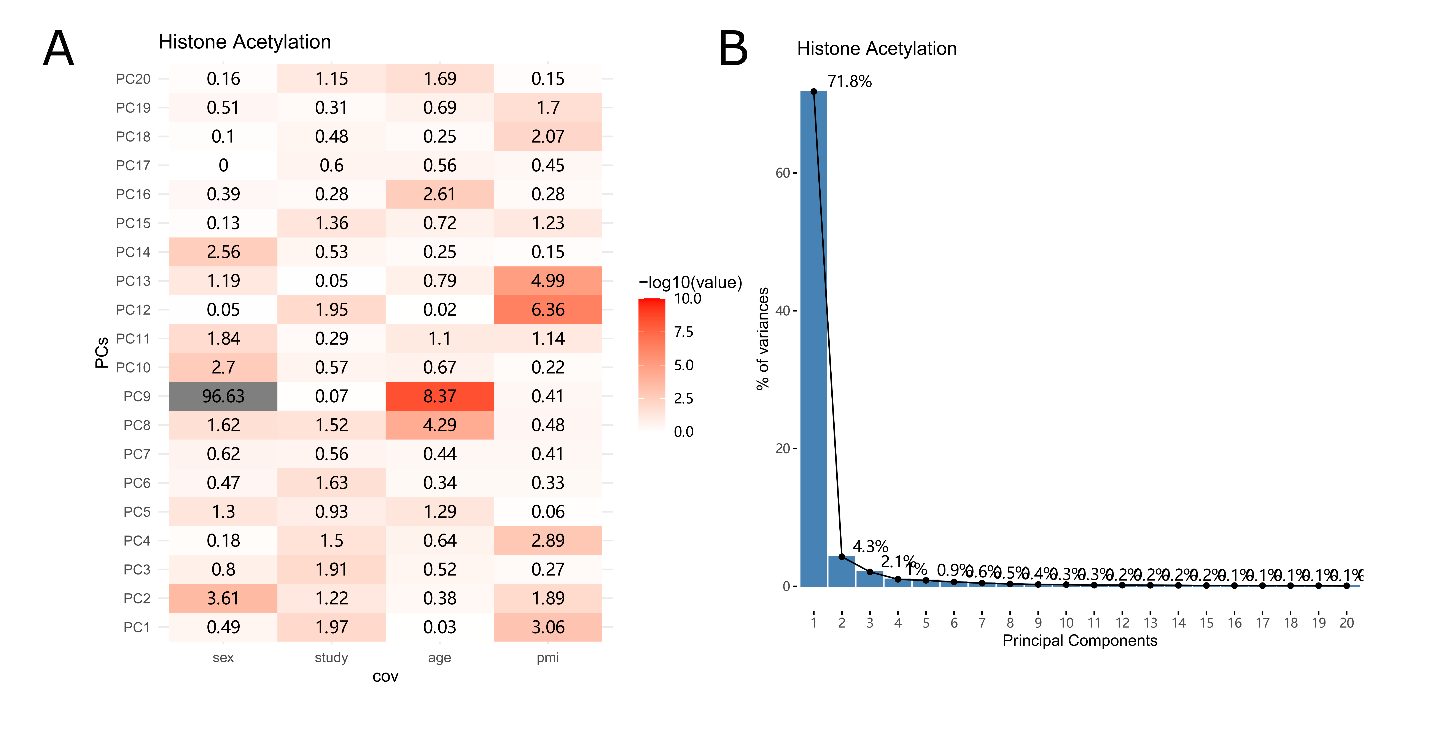

###### *Supplementary Figure 4. Top 20 Principal Components (PCs) from histone acetylation data A) Significances of association between top 20 PCs and four covariates derived from histone acetylation data, including sex, study, age of death and PMI, derived from Omnibus F-tests for linear regression models B) Scree plots illustrating variances explained by top 20 PCs for histone acetylation data.*

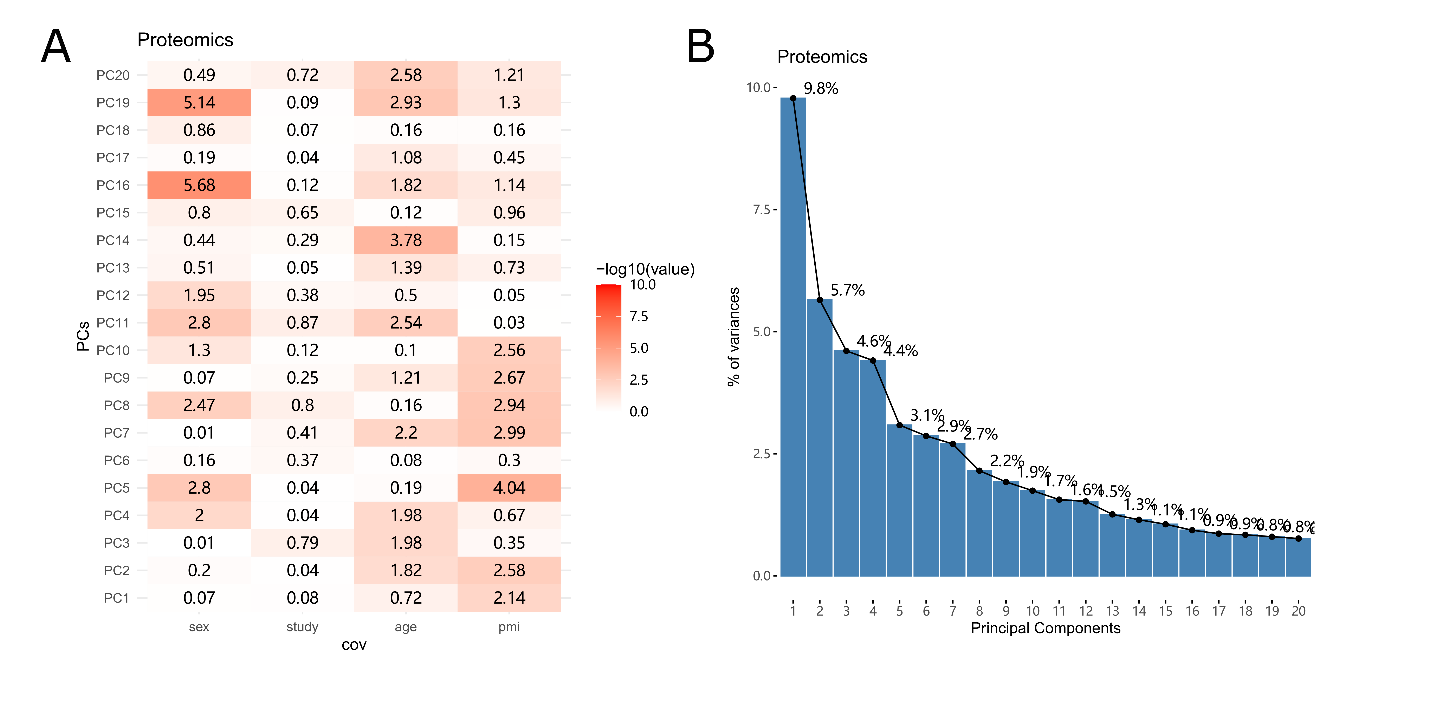

###### *Supplementary Figure 5. Top 20 Principal Components (PCs) from proteomic data A) Significances of association between top 20 PCs and four covariates derived from proteomic data, including sex, study, age of death and pmi, derived from Omnibus F-tests for linear regression models B) Scree plots illustrating variances explained by top 20 PCs for proteomic data.*

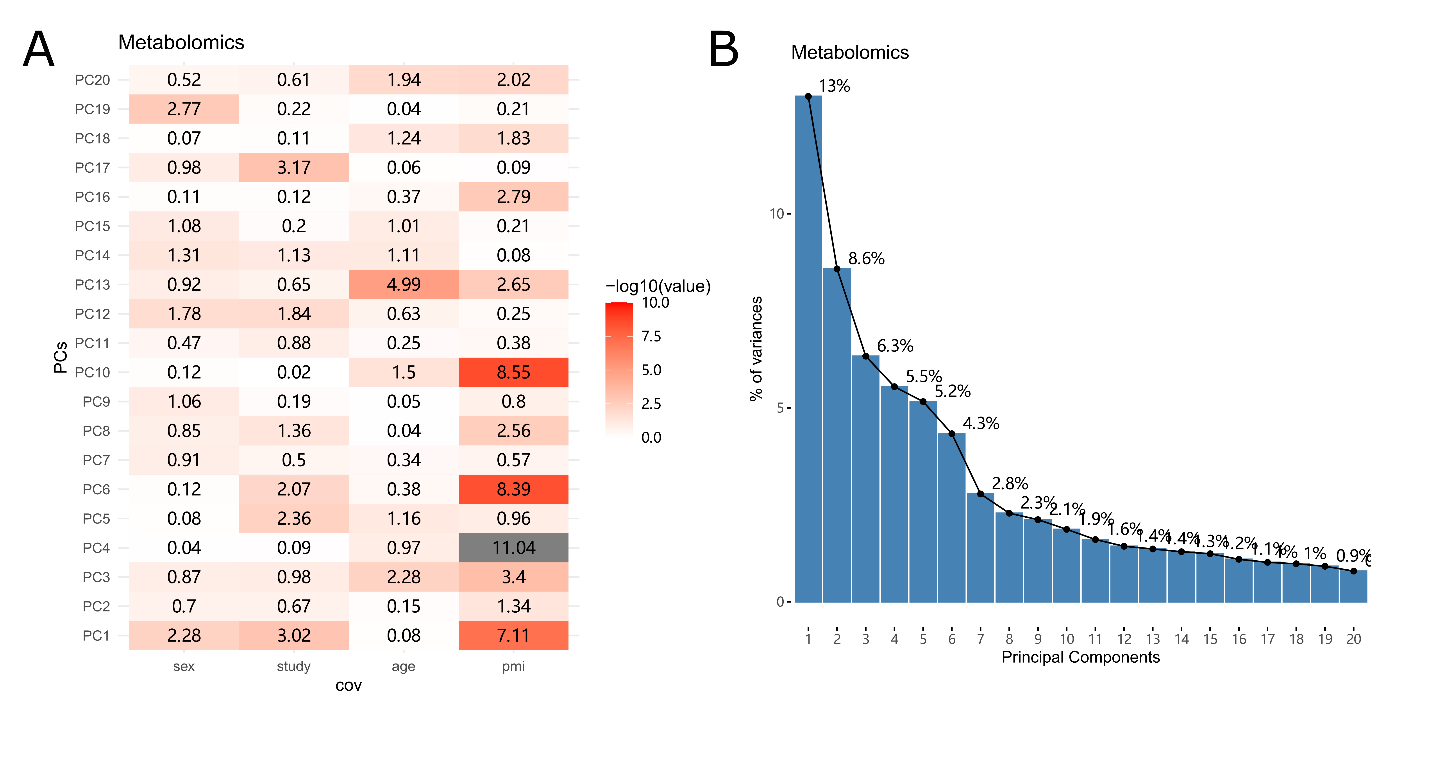

###### *Supplementary Figure 6. Top 20 Principal Components (PCs) from metabolomic data A) Significances of association between top 20 PCs and four covariates derived from metabolomic data, including sex, study, age of death and pmi, derived from Omnibus F-tests for linear regression models B) Scree plots illustrating variances explained by top 20 PCs for metabolomic data.*

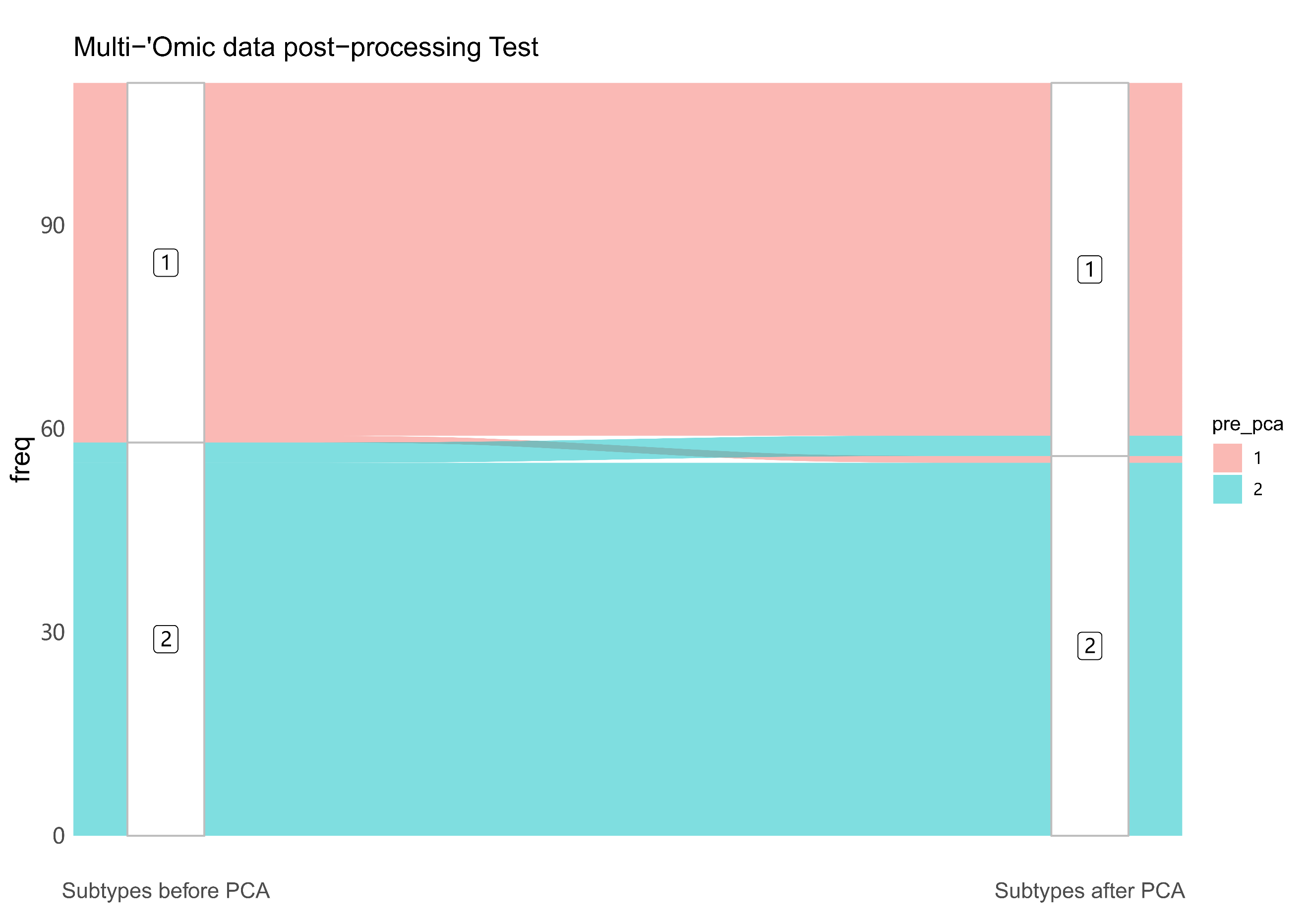

###### *Supplementary Figure 7. Testing the similarity between subgroups with and without post-processing (PCA) Alluvial plot comparing the subtype results for 5-modality integration on individual level before and after post-processing. The optimal subtype number was assigned as 2 identified unanimously agreed by eigen-gaps and dip-test statistics.*

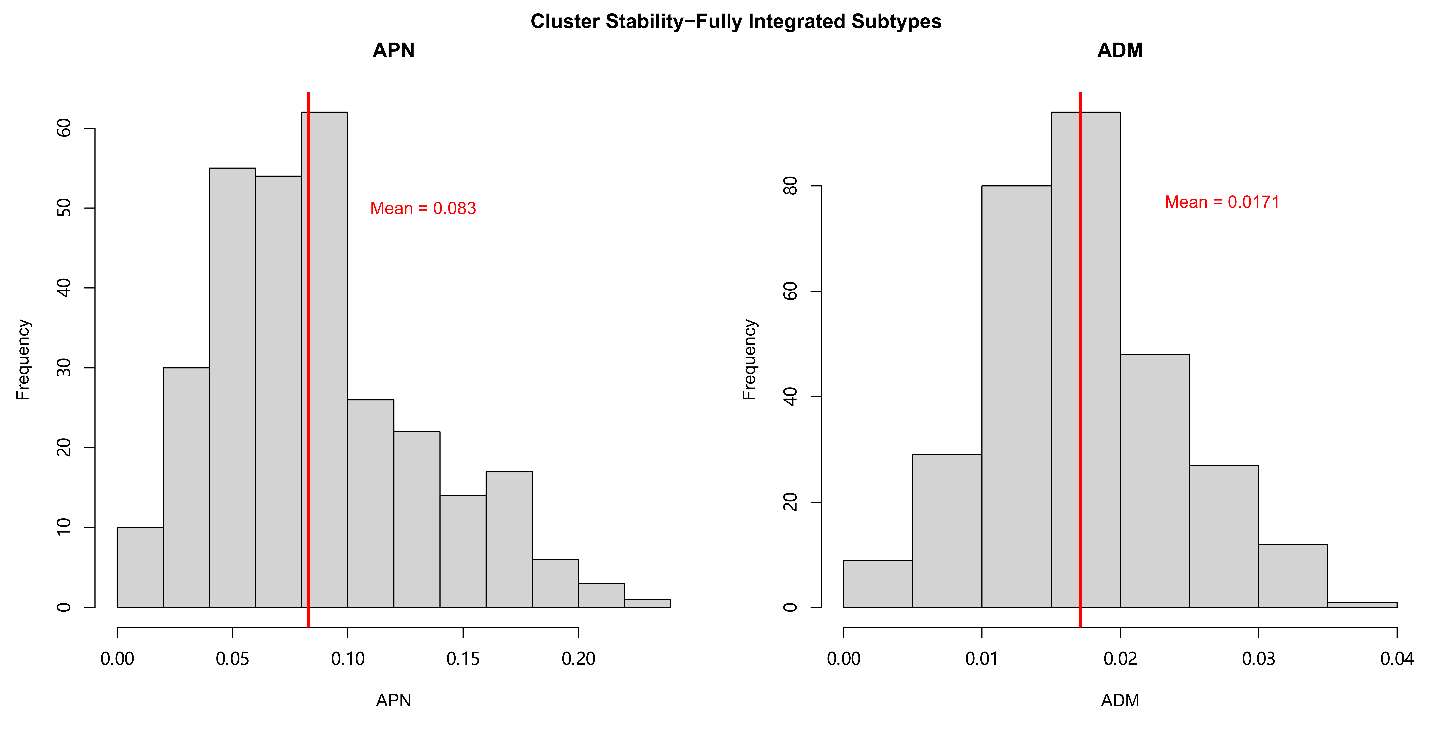

###### *Supplementary Figure 8. Cluster stability analysis performed on fully integrated molecular subtypes Histograms for the distribution of both APN and ADM generated from 300 random sub-samples for the fully integrated subtypes.*

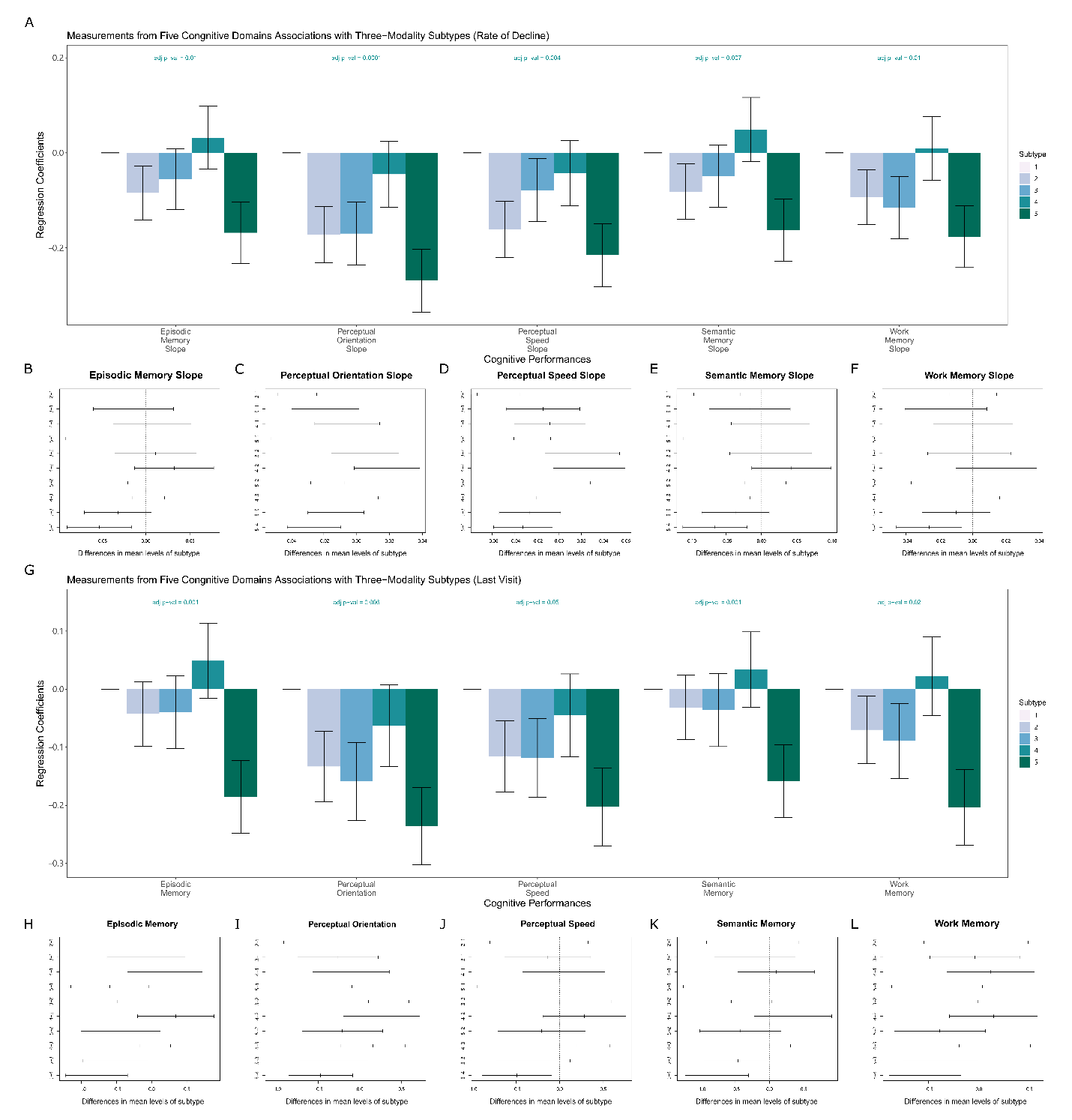
*Supplementary Figure 9. Molecular subtypes derived from histone acetylation, DNA methylation and RNAseq were tested against cognitive performance at last visit and rate of cognitive performance decline in 5 domains A) Consensus associations of 3-modal integrated subtypes and rate of cognitive performance decline from five domains. Y-axis shows standardized beta coefficients estimated from linear regression, where subtype 1 was used as the baseline category. Positive (negative) standardized beta coefficients indicate a superior (inferior) performance. Error bars indicate one standard deviation value for estimated beta coefficients. B-F): Difference in mean value of rate of cognitive performance decline from each of the five domains at last visit between subtypes by Tukey’s HSD. G) Consensus associations of 3-modal integrated subtypes and cognitive performance at last visit from five domains. Association method used was the same as A. H-L): Difference in mean value of cognitive performance from each of the five domains at last visit between subtypes by Tukey’s HSD.*

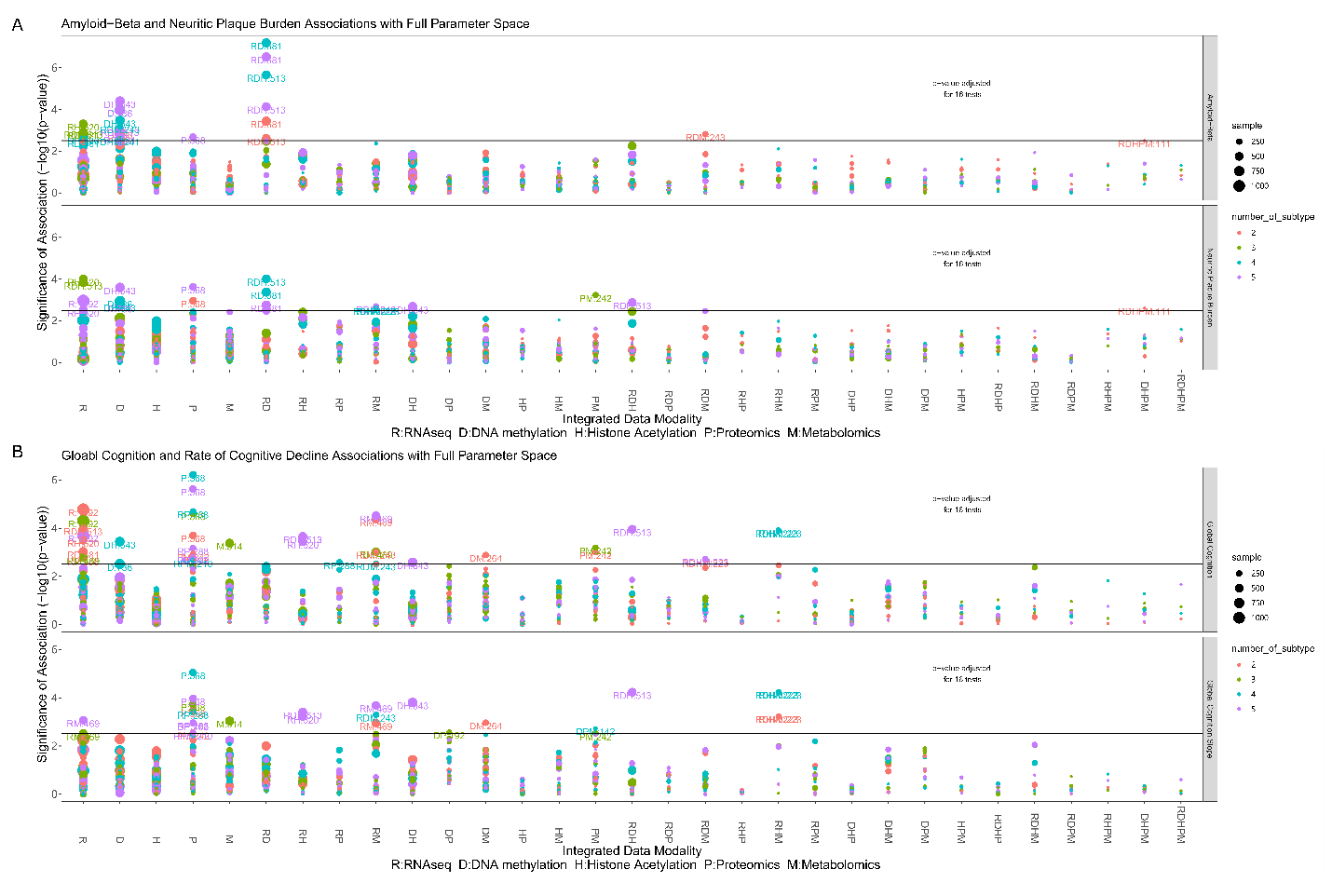

###### *Supplementary Figure 10. Full parameter search space tested on cognitive measurements and neuropathologies. A)* *Associations of subtypes identified from the full parameter search space and global cognition at last visit and rate of cognitive decline (844 unique combinations and 13,504 corresponding tests) by linear regression models.* *Y-axis shows significance of association (-log_10_ transformed raw p-values). The black horizontal line illustrates an unadjusted p-value threshold at 0.05, whereas the purple horizontal line demonstrates Bonferroni-adjusted p-value thresholds for 16 tests (p_raw_=3.1x10^-3^). X-axis labels represent integrated data modalities used for SNF (R: RNAseq, D: DNA methylation, H: Histone acetylation, P: Proteomics, M: Metabolomics, RD: integrating RNAseq and DNA methylation etc.). Labels on plotted data illustrate the sample size and corresponding sample size identified by data modalities, whereas the sample size is represented by the size of plotted data points. B) Associations of subtypes identified from the full parameter search space and amyloid-beta and neuritic plaque burden.*

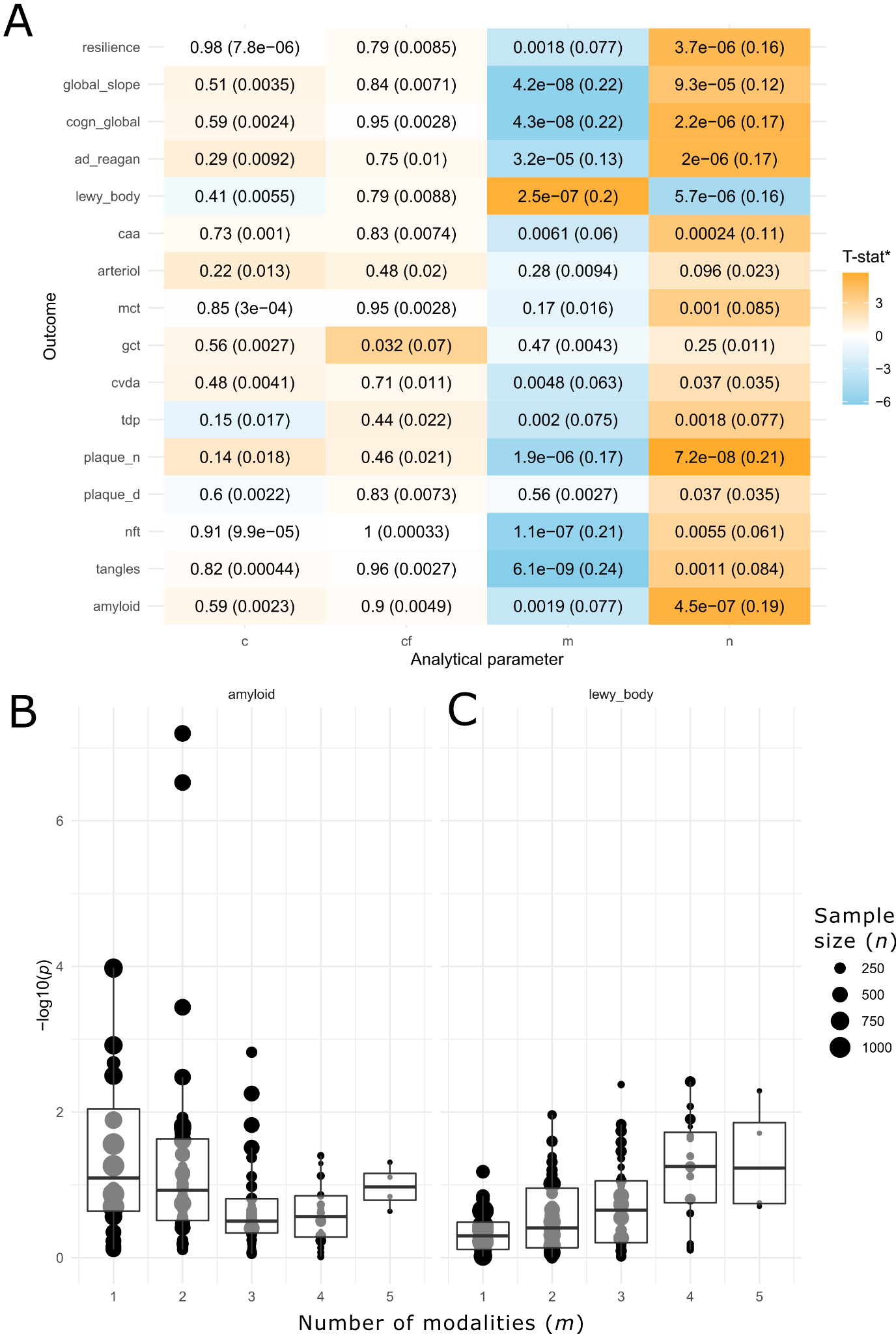

### ***Supplementary Figure 11.*** *Results of meta-regression of external validity for subtyping solutions across pipeline parameters.A)* *Heatmap showing T-statistics (*F-statistics are provided for the cf parameter analysis, as it was treated as a categorical factor and therefore analyzed with ANOVA), uncorrected two-sided p-values and model variance explained (r^2^, in brackets) for the effect of each parameter (c, cf, m, and n) on the association significance (-log(p)) of resulting subtypes with each neuropathological and cognitive variable. B-C) Boxplots showing the significance of cluster membership association ships with amyloid and Lewy body neuropathology across values of m (number of modalities, x-axis) and n (sample size; dot size). amyloid=total beta amyloid; arteriol=arteriolosclerosis; caa=cerebral amyloid angiopathy; cvda=cerebral atherosclerosis; gct=gross cerebral infarcts; mct=cerebral microinfarcts; nft=neurofibrillary tangles count; plaq_d=diffuse plaque count; plaq_n=neuritic plaque count; tangles=paired helical filament tau.*

### Supplementary Tables

###### *Supplementary Table 1. Table summarizing Normalized Mutual Information (NMI) values between fused networks from all 5 data modalities and each data modalities alone*

|  | Fused Network | RNASeq | DNA-Methylation | Histone Acetylation | Proteomic | Metabolomic |
| --- | --- | --- | --- | --- | --- | --- |
| Fused Network | 1 | **0.15** | **0.18** | **0.38** | 0.04 | 0.05 |
| RNASeq |  | 1 | 0.012 | 0.009 | 0.007 | 0.007 |
| DNA-Methylation |  |  | 1 | 0.05 | 0.006 | 0.002 |
| Histone Acetylation |  |  |  | 1 | 0.06 | 0.003 |
| Proteomic |  |  |  |  | 1 | 0.002 |
| Metabolomic |  |  |  |  |  | 1 |

*Note: NMI ranges between 0 and 1, higher NMI values represents more similarities between the two data modalities*

###### *Supplementary Table 2. Table summarizing top 10 features from top contributor modalities for five-modal integrated subtypes identified by ANOVA*

| Data Modalities | Feature Name | Genomic Region/Gene |
| --- | --- | --- |
| RNASeq | ENSG00000102699 | PARP4 |
|  | ENSG00000204052 | LRRC73 |
|  | ENSG00000164938 | TP53INP1 |
|  | ENSG00000185650 | ZFP36L1 |
|  | ENSG00000099960 | SLC7A4 |
|  | ENSG00000138028 | CGREF1 |
|  | ENSG00000166448 | TMEM130 |
|  | ENSG00000067141 | NEO1 |
|  | ENSG00000103335 | PIEZO1 |
|  | ENSG00000168539 | CHRM1 |
| DNA-methylation | cg16783349 | chr10:130,854,343 |
|  | cg07860645 | DAAM2 |
|  | cg16357930 | PLXNB1 |
|  | cg17367884 | TBC1D16 |
|  | cg00501869 | PTPRN2 |
|  | cg06930722 | CHST11 |
|  | cg01005441 | ARHGEF10 |
|  | cg14040679 | chr9:98,811,442 |
|  | cg15355952 | SLC1A3 |
|  | cg07652350 | chr11:616,009 |
| H3K9 Histone Acetylation | peak19236 | NDST3 |
|  | peak21593 | ELOVL4 |
|  | peak6042 | SYT1 |
|  | peak20453 | NDFIP1 |
|  | peak20236 | CHSY3 |
|  | peak11660 | OSBPL1A |
|  | peak15109 | PECR, TMEM169 |
|  | peak18276 | IL12A-AS1, C3orf80 |
|  | peak18140 | CLSTN2 |
|  | peak4863 | MTMR2 |

###### *Supplementary Table 3. Table summarizing top 10 features from each modalities for 3-modal integrated subtypes identified by ANOVA tests*

| Data Modalities | Feature Name | Genomic Region/Gene |
| --- | --- | --- |
| RNASeq | ENSG00000145882 | PCYOX1L |
|  | ENSG00000254561 | NECTIN1 |
|  | ENSG00000198743 | SLC5A3 |
|  | ENSG00000163605 | PPP4R2 |
|  | ENSG00000186298 | PPP1CC |
|  | ENSG00000179477 | ALOX12B |
|  | ENSG00000100577 | GSTZ1 |
|  | ENSG00000132155 | RAF1 |
|  | ENSG00000100360 | IFT27 |
|  | ENSG00000169762 | TAPT1 |
| DNA-methylation | cg26878318 | chr5:10,308,785 |
|  | cg21512370 | RB1, LPAR6 |
|  | cg24769903 | RP11-83B20.10 |
|  | cg01882222 | chr5:172,407,583 |
|  | cg09357831 | chr7:100,214,622 |
|  | cg10088941 | chr6:43,608,807 |
|  | cg02489379 | AC010149.4 |
|  | cg15590863 | FAM185A |
|  | cg10616859 | BRAT1 |
|  | cg04276288 | CTAGE5 |
| H3K9 Histone Acetylation | peak7234 | ZNF219, TMEM253 |
|  | peak12821 | LSM14A |
|  | peak10522 | PSMD11, CDK5R1, MYO1D, RP11-466A19.1 |
|  | peak10347 | ALDH3A2, CTD-2104P17.2 |
|  | peak18869 | APBB2 |
|  | peak20390 | LINC01024, PURA |
|  | peak1727 | MEF2D,Y_RNA |
|  | peak16650 | RHBDD3, EWSR1 |
|  | peak10319 | MYO15A, ALKBH5 |
|  | peak10169 | ZBTB4, SLC35G6 |

###### *Supplementary Table 4. Full dataset of summary statistics from sensitivity analysis*

###### *Column description:*

###### *“limit_by”: Data modalities combinations which limited the sample size*

*“clust_num”: Assigned cluster number prior to perform spectral clustering*

*“opt_est”: The optimal cluster solution estimated by Eigen-gap and rotation cost method, smaller cluster number is recorded when two methods disagree.*

*“snf_type”: Data modalities used for SNF*

*“sample”: Sample size*

*“response”: Corresponding response variable: either neuropathologies or cognitive measures*

*“p”: Raw p-value from F-test*

*“n_modes”: Number of data modalities used in SNF*

*Full table found in available in a separate file*
